# CASTER-DTA: Equivariant Graph Neural Networks for Predicting Drug-Target Affinity

**DOI:** 10.1101/2024.11.25.625281

**Authors:** Rachit Kumar, Joseph D. Romano, Marylyn D. Ritchie

**Affiliations:** Medical Scientist Training Program, Perelman School of Medicine, University of Pennsylvania, USA; Genomics and Computational Biology Graduate Group, Perelman School of Medicine, University of Pennsylvania, USA; Institute for Biomedical Informatics, Perelman School of Medicine, University of Pennsylvania, USA; Center of Excellence in Environmental Toxicology, Perelman School of Medicine, University of Pennsylvania, USA; Department of Biostatistics, Epidemiology, and Informatics, Perelman School of Medicine, University of Pennsylvania, USA; Department of Genetics, Perelman School of Medicine, University of Pennsylvania, USA

**Keywords:** graph neural networks, structural biology, deep learning, protein representation learning

## Abstract

Accurately determining the binding affinity of a ligand with a protein is important for drug design, development, and screening. With the advent of accessible protein structure prediction methods such as AlphaFold, predicted protein 3D structures are readily available; however, methods for predicting binding affinity currently do not take full advantage of 3D protein information. Here, we present CASTER-DTA (Cross-Attention with Structural Target Equivariant Representations for Drug-Target Affinity), which uses an equivariant graph neural network to learn more robust protein representations alongside a standard graph neural network to learn molecular representations to predict drug-target affinity. We augment these representations by incorporating an attention-based mechanism between protein residues and drug atoms to improve interpretability. We show that CASTER-DTA represents a state-of-the-art improvement on multiple benchmarks for predicting drug-target affinity and that it generates novel insights for several related tasks. We then apply CASTER-DTA to create a large resource of the binding affinities of every FDA-approved drug against every protein in the human proteome and make these predictions freely available for download. We also make available a web server for researchers to apply a pretrained CASTER-DTA model for predicting binding affinities between arbitrary proteins and drugs.

**Key Messages:**

- CASTER-DTA is a novel architecture that makes use of equivariant graph neural networks to predict drug-target affinity, enabling rapid and interpretable predictions about how proteins and molecules bind to each other.
- Equivariant graph neural networks like those used in CASTER-DTA allow for more robust usage of protein 3D structural information and improve performance over non-equivariant approaches.
- CASTER-DTA includes a cross-attention layer to update protein residue and molecule atom embeddings that allows for interpretability about which residues are involved in the prediction.
- We can use this architecture for a variety of downstream tasks, and we have used CASTER-DTA to generate a resource of predicted binding affinities for every FDA-approved drug against every protein in the human proteome.

## Introduction

Determining drug-target affinity (DTA) is a critical component of the drug design and discovery process, serving as a metric of which drugs (ligands/molecules) may be high-priority for further testing in terms of binding to a target (protein)[1]. While experimental affinity determination is the gold standard, it is often not feasible or practical to perform experiments to determine affinity for what may be millions or billions of possible drug candidates for a given target. To address this, various computational methods have been developed for the purpose of predicting the binding affinity of an arbitrary protein and an arbitrary ligand to help prioritize or rank drugs for further exploration.

Some state-of-the-art methods, such as DeepDTA and AttentionDTA, use primarily sequence information, using the protein amino-acid sequence and the molecule Simplified Molecular Input Line Entry System (SMILES) string [2, 3]. Other methods, such as GraphDTA and DGraphDTA, make use of graph neural networks (GNNs) to learn protein or molecular representations[4, 5].

To date, few methods have made use of the underlying geometric information contained within the entire protein 3D structure such as bond angles or direction vectors for predicting binding affinity. Furthermore, to our knowledge, no methods have made use of SE(3)-equivariant graph neural networks specifically to use these protein-level features in a rigorous way for the purposes of drug-target affinity prediction.

We present in this paper a method, **C**ross-**A**ttention with **S**tructural **T**arget **E**quivariant **R**epresentations for **D**rug-**T**arget **A**ffinity (CASTER-DTA), inspired by work done in other paradigms[6, 7, 8], that makes use of SE(3)-equivariant graph neural networks in the form of Geometric Vector Perceptron (GVP)-GNNs to incorporate this 3D backbone information at the protein level alongside a standard graph neural network to process molecular information augmented by cross-attention to improve drug-target affinity prediction. We show that CASTER-DTA outperforms other methods in this paradigm and highlight its utility on a variety of downstream tasks.

A graphical abstract of the datasets, training process, architecture, and downstream analyses that we performed in this paper can be found in **Figure 1**.

**Fig. 1.**
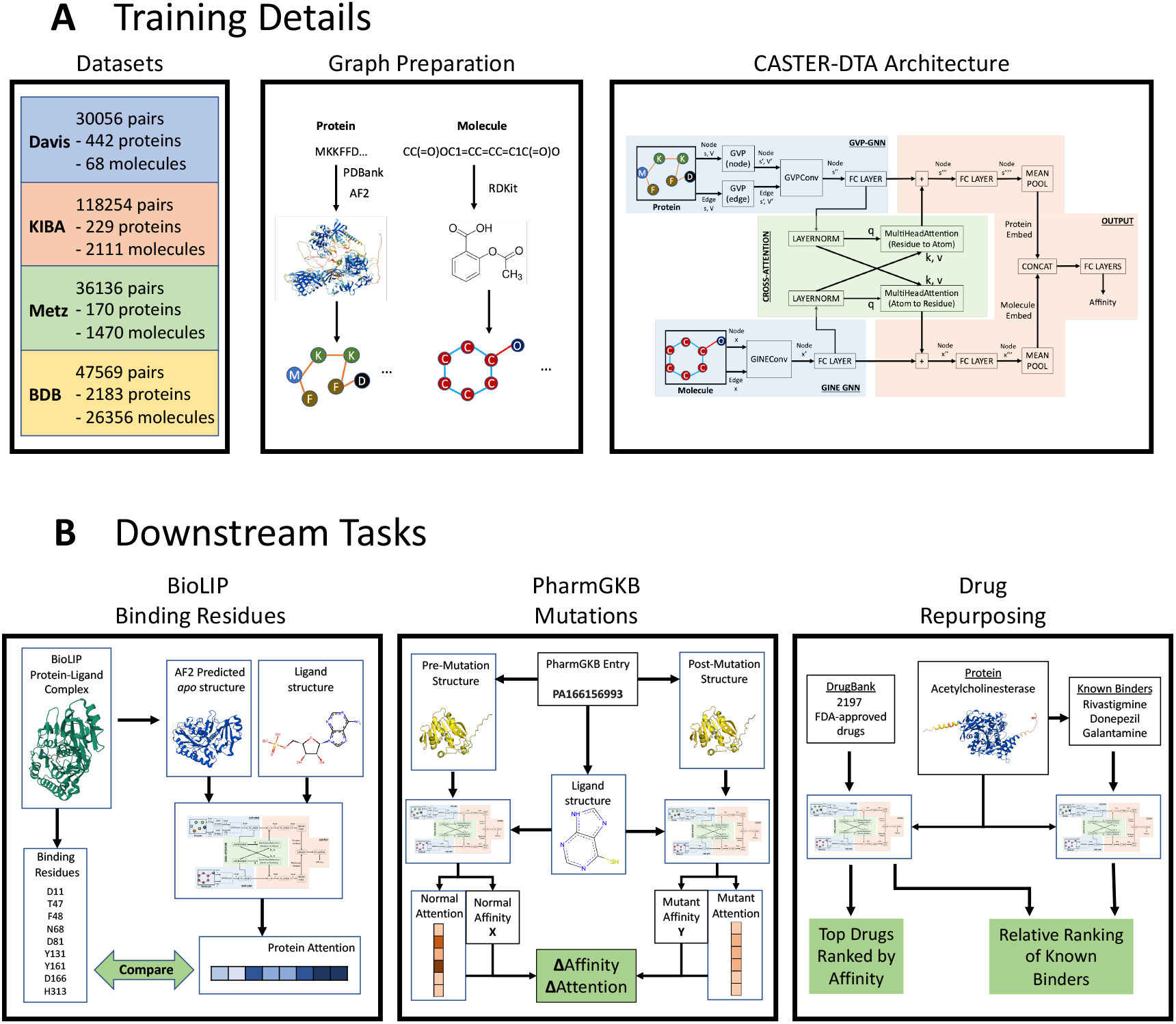
A visual overview of multiple aspects of this paper. (A) A description of the datasets, graph preparation, and architecture used in the training process of this paper. (B) A description of each of the downstream tasks performed in this paper, including BioLIP binding residue assessment, PharmGKB mutation analysis, and an analysis of drug repurposing for known proteins and drugs. The light green boxes or arrows represent the endpoint of each analysis for validation of CASTER-DTA.

## Methods

### Dataset Details and Preparation

To evaluate our model, we used four established datasets for the purposes of predicting drug-target affinity: the Davis dataset [9], the KIBA dataset [10], the Metz dataset [11], and the BindingDB (*K*_*d*_) dataset[12] (with specific filters, described below). Details on the number of protein-molecule pairs, unique proteins, and unique molecules in each dataset are provided in **Table 1**, along with information about the target values for each dataset.

**Table 1.** Binding Affinity Dataset Information

| Dataset | Unique Proteins | Unique Molecules | Total Pairs | Target Values |
| --- | --- | --- | --- | --- |
| Davis | 442 | 68 | 30056 | $pK_d^*$ |
| KIBA | 229 | 2111 | 118254 | KI score |
| Metz | 170 | 1470 | 36136 | $pK_d$ |
| BindingDB | 2183 | 26356 | 47569 | $pK_d^*$ |

\*Converted from provided $K_d$ value using **Equation 1**.

We use then BindingDB December 2024 release, and we only use pairs with a defined, exact *K*_*d*_ value (that is, not binned such as “*<*0.5 *µ*M”) and where the ligand produces a valid molecule in RDKit. We also filter to pairs where the protein has a sequence length of less than 3000 residues as several of the benchmark methods clip sequences below this length, though CASTER-DTA can operate on longer proteins.

All datasets provide amino-acid sequences for each protein and SMILES strings for each molecule. The Davis and KIBA datasets have isomeric SMILES available, while the Metz and BindingDB datasets provide canonical SMILES.

For KIBA, the target is provided as an integrated bioactivity score that integrates *K*_*d*_, *K*_*i*_, and IC50 values from each protein-ligand pair. For Davis and BindingDB, the target is the *K*_*d*_ value representing the binding affinity between each protein-molecule pair. *K*_*d*_ values are converted to scaled *pK*_*d*_values by performing the transformation in **Equation 1**.

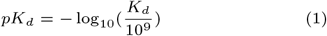

For Metz, the target is given as a scaled *pK*_*d*_ value, the same as the post-scaled Davis and BindingDB datasets.

We acquired ligand-free (*apo*) structures for every protein in each dataset in the following manner, using the first method that yielded a result:

1. Search the Protein DataBank (PDBank) [13] for experimentally-determined structures with 100% sequence identity, filtering to only *apo* structures. If multiple resolved structures exist, the structure with the best resolution was used.
2. Search the AlphaFold2 database (co-hosted on PDBank) for pre-folded structures with 100% sequence identity [14]. If multiple matches existed, the result with the highest pLDDT was used.
3. Use a local version of AlphaFold2 [15] (ColabFold, using colabfold batch [16]) to fold any remaining proteins using the sequences provided, with settings described in **Appendix A**.

### Protein Graph Representations

We construct protein graphs from the *apo* protein structures, inspired by previous work into using equivariant graph neural networks for other protein-related tasks; in particular, we took inspiration from the PocketMiner GNN for predicting cryptic pocket opening within proteins [8] and the original GVP-GNN papers and its extension [6, 7]; however, instead of connecting to the 30 nearest neighbors as in those approaches, we connect each residue to every residue within a 4 Angstrom distance (including a residue to itself).

For protein node (residue) and edge features, we include both scalar and vector-based features as inspired by PocketMiner and other work, as described in **Appendix B.1**. We concatenate to the node scalar features a one-hot-encoded representation of the amino acid identity with 20 unique amino acids and an extra class to represent nonstandard amino acids if they are present in the dataset.

### Molecular Graph Representations

We construct 2D molecular graphs from SMILES strings by using RDKit [17], where each node represents an atom and each edge represents a bond between atoms or a self-loop from an atom to itself. For molecule node (atom) and edge (bond) features, we include a variety of features for atomic representations based on prior work as described in **Appendix B.2**.

Additionally, we concatenate one-hot encoded atom types, including one of 10 unique types representing hydrogen, carbon, nitrogen, oxygen, fluorine, phosphorus, sulfur, chlorine, bromine, iodine, and other, respectively, where “other” captures all other atom types. Additionally, a one-hot-encoded representation of whether an edge represents a single, double, triple, or aromatic bond or self-loop is concatenated to the edge features as a representation of the bond type.

### Architecture

CASTER-DTA includes two graph neural networks that produce learned representations at the node level. Protein graphs are processed by Geometric Vector Perceptron (GVP) and GVPConv layers which are SE(3)-equivariant, as proposed by Jing et al.[6, 7]. Initial GVPs are used to create richer representations of the node and edge features before processing by GVPConv layers to update the node-level (residue) representations. Molecule graphs are processed by GINEConv layers, which represent an edge-enhanced version of the Graph Isomorphism Network (GIN) as described by Xu et al. and Hu et al. [18, 19]. Each GINEConv layer updates the node-level (atom) representations.

The node-level representations for proteins and molecules are subsequently processed by a linear layer and then used as queries, keys, and values in a cross-attention paradigm, where each cross-attention block updates either the protein or molecule embeddings based on the attention weights computed for residues attending to atoms or vice-versa. These updated node embeddings are processed by another linear layer and are each then pooled by taking the mean over all nodes to create a singular graph embedding for proteins and molecules; these embeddings are subsequently concatenated and passed into several fully-connected (FC) layers for the final regression prediction. A comprehensive visual summary of this architecture can be seen in **Figure C1**.

In evaluating this model, we assessed different combinations based on the number of convolutional layers for each of the protein and molecule GNNs, with either 1 or 2 layers being selected for each of the protein or molecule GNNs. This totals 4 combinations, defined as CASTER-DTA(1,1), CASTER-DTA(1,2), CASTER-DTA(2,1), and CASTER-DTA(2,2), where the number in the parentheses (X,Y) refers to the number of protein convolutions (X) and molecule convolutions (Y), respectively.

### Training and Evaluation

We split each dataset into 70% training, 15% validation, and 15% testing splits, doing so multiple times to evaluate the model’s performance and consistency.

When training CASTER-DTA, the output values for each dataset are standardized to have a mean of 0 and a standard deviation of 1 during training. The model results are subsequently unscaled for the purposes of reporting and evaluation, and all metrics below for CASTER-DTA are computed on the unscaled outputs.

We trained CASTER-DTA on the training splits for 2000 epochs using an Adam optimizer with a base learning rate of 1e-4 on a Mean-Squared Error (MSE) loss function. We multiply the learning rate by 0.8 after 50 epochs of no observed improvement and end training early if the validation loss does not improve for 200 epochs. In most cases across architectures, CASTER-DTA terminated around 1000 epochs.

Due to variations in the size of protein and molecule graphs, memory usage can fluctuate dramatically from batch to batch. To address this and stabilize training, for the Davis and Metz datasets, we employ a dynamic mini-batching paradigm wherein we limit the total size of a batch to 16 million residue-atom pairs with a maximum of 128 protein-molecule pairs in each batch. For the KIBA dataset, we limit to 8 million residue-atom and 64 protein-molecule pairs, and for the BindingDB dataset, we limit to 4 million residue-atom and 32 protein-molecule pairs. This difference is due to different memory requirements due to larger proteins being present in the KIBA and BindingDB dataset. All models were trained on NVIDIA RTX 2080Ti (12GB) or NVIDIA A30 (24GB) GPUs. The A30s were used as-needed to train when datasets or models were too large to fit on the smaller 12GB GPUs, necessary for some of the ablation studies that we performed.

We used the model with the best validation loss during each of the above training cycles for downstream evaluation on the test set, doing so across five different training/validation/testing splits (defined by seeds). To compare, we also trained and evaluated several preexisting state-of-the-art models on the same train/validation/test dataset splits, including DeepDTA, GraphDTA, DGraphDTA, DeepGLSTM, and AttentionDTA. All hyperparameters for these models were replicated from their respective papers for both datasets, and these hyperparameters are described in **AppendixC**. DGraphDTA could only be evaluated on Davis and KIBA due to its reliance on MSA-related features that were only provided for these two datasets.

### Ablation Studies

We performed an ablation study where we replaced the protein GNN with a graph attention network (GATv2) convolutional layer [20, 21] instead, as GAT has been used in prior work for protein graph processing [5]. We also assessed a variety of molecular GNN architectures to determine which molecular representation worked best for CASTER-DTA, including GIN [18], GINE [19], GATv2 [20, 21], and AttentiveFP [22]. We performed these tests on the same five seeds of the Davis dataset with the same training/testing paradigm as described above using a model with two protein GNN convolutions and two molecule GNN convolutions (analogous to CASTER-DTA(2,2)).

We also assessed how different edge definitions impacted the model, testing Angstrom distance thresholds of 4, 6, and 8 Angstroms. We also assessed using the 30 nearest neighbors. We performed this ablation study using the base architecture of CASTER-DTA(2,2) on the Davis dataset.

### Interpretability Analyses and Validation

To assess the interpretability of CASTER-DTA, we used the best model trained on the BindingDB dataset, in this case one of the CASTER-DTA(2,2) models, to predict the binding affinity of various proteins. We used the model trained on the BindingDB dataset as such a model most likely has the highest generalizability, given the high number of diverse proteins and molecules in that dataset (**Table 1**).

CASTER-DTA generates attention matrices from its cross-attention module, which we use to assess which residues of the protein are most important for the binding of the drug by computing the average attention score for each residue over all molecule atoms.

The base expected attention score depends on the length of the protein; that is, a protein of length 1000 would have an expected attention score of 1/1000, or 0.001, for each residue. To address this, we compute a length-scaled ratio where a value of 0 means that the residue has been assigned an attention score equal to the expected score, a negative value means that the residue has been attended to less, and a positive value means that the residue has been attended to more. This process is described in **Equation 2**.

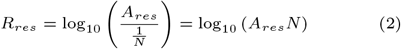

We assessed proteins where mutations were known to impact the protein’s binding to certain drugs or where the binding site of particular ligands was known or predicted to determine if the distribution of attention scores indicated an increased focus on residues involved in the mutation or binding site.

### BioLIP Binding Residue Assessment

To assess whether CASTER-DTA’s attention mechanism highlights residues that are ligand-binding, we use the BioLIP dataset [23]. BioLIP provides information on putative ligand-binding residues for each protein-ligand interaction in its database determined through intermolecular contacts. We use all protein-ligand pairs where: (1) the ligand has a SMILES that can be processed by RDKit into a valid graph; (2) the protein has an existing structure in the preexisting AlphaFold2 human database; and (3) the protein or ligand do not have unknown/missing atom or residue types. This results in 936 unique protein-ligand pairs, comprised of 383 unique ligands and 754 unique proteins.

We subset each protein to the sequence present in the BioLIP database and run CASTER-DTA on each protein-ligand complex, extracting the attention scores. Then, to assess whether CASTER-DTA “focuses” on binding residues, we compute for each complex two values: the average attention score over the binding residues and the average attention score over the nonbinding residues. We then perform a paired t-test between these values over all complexes to test the hypothesis of whether the attention values are different between the two groups.

### PharmGKB Mutation Impact Assessment

To assess whether CASTER-DTA’s attention mechanism preferentially highlights mutated protein residues, we use PharmGKB’s manually curated list of Very Important Pharmacogenes (VIPs), a list that highlights the most clinically-actionable mutations, to find a variety of mutations that affect drug metabolism[24]. We select four protein-coding mutations associated with a change in protein sequence where protein structure may change as a result of the mutation.

Using CASTER-DTA, we determine the change in predicted binding affinity between the nonmutant and mutant versions of the protein and compare to experimental observ; we also assess whether the introduction of a mutation at specific residues leads to differences in attention at those residues specifically. The mutations that we assessed in this manner were rs61742245, rs1800462, rs4149056, and rs2306283, and information on the gene that the mutation maps to, the drug that the mutation is purported to affect, the NCBI ID of the protein that the mutation affects, and the amino acid change in the protein that the mutation causes can be found in **Table 2**.

**Table 2.**
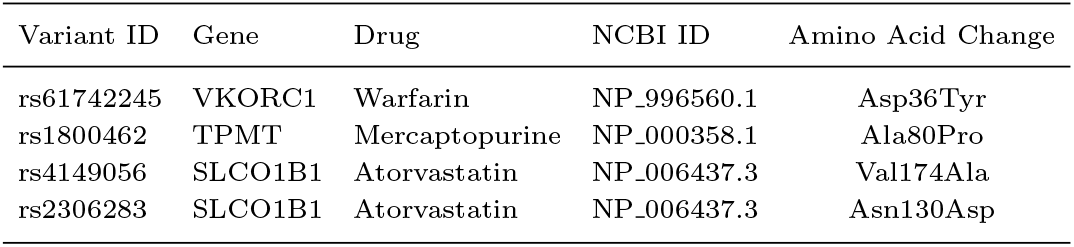
PharmGKB Mutations Assessed

| Variant ID | Gene | Drug | NCBI ID | Amino Acid Change |
| --- | --- | --- | --- | --- |
| rs61742245 | VKORC1 | Warfarin | NP_996560.1 | Asp36Tyr |
| rs1800462 | TPMT | Mercaptopurine | NP_000358.1 | Ala80Pro |
| rs4149056 | SLCO1B1 | Atorvastatin | NP_006437.3 | Val174Ala |
| rs2306283 | SLCO1B1 | Atorvastatin | NP_006437.3 | Asn130Asp |

### Drug Repurposing

For both drug repurposing tasks, we also used the best CASTER-DTA(2,2) model trained on the BindingDB dataset for the same reasons as above.

#### Case Studies for Validation

For validation, we use two relevant case studies in Alzheimer’s disease (AD) and H5N1 bird flu where strong evidence exists that certain proteins may be involved in either the pathogenesis or symptoms of disease and where they have a public health relevance. These proteins include: Amyloid-beta peptide, Amyloid precursor protein, Acetylcholinesterase, and Tau for Alzheimer’s disease; and H5 Hemagglutinin, H5N1 Nucleoprotein, and N1 Neuraminidase for H5N1 bird flu. The set of proteins assessed are repeated and extended in **Table E1** with additional information on the Protein DataBank ID associated with the structures that were used for their 3D structures.

We get a list of all FDA-approved drugs from DrugBank, filtering out withdrawn drugs or drugs with invalid SMILES. This results in 2197 drugs, and we assess the binding affinity of all of these drugs against the above proteins. We report the top three drugs for each protein and whether prior research shows that they may have utility against that protein or if they are novel.

To determine how our model behaves for known existing drugs, we also assess the predicted *pK*_*d*_ and z-scores of existing FDA-approved drugs known to bind to Acetylcholinesterase and Neuraminidase. For Acetylcholinesterase, these drugs are rivastigmine, donepezil, and galantamine. For Neuraminidase, we assessed oseltamivir, zanamivir, and peramivir.

#### Proteome-Wide Assessment

To provide a rich resource for future researchers without requiring them to set up and run CASTER-DTA themselves, we provide computational predictions for the binding affinity of all FDA-approved drugs against all proteins currently recorded in the human proteome.

For this prediction, we acquire all AlphaFold2 structure predictions for the human reference proteome (UP000005640), comprising 23,391 predicted structures [14] and acquire all SMILES strings for FDA-approved drugs in the ZINC2020 database, comprising 1615 unique drugs [25]. We used the ZINC20 database as the source for the FDA-approved drug data to ensure that we could distribute the resulting predictions without restriction. We use CASTER-DTA to predict binding affinities for all resulting 37,776,465 pairs and make these predictions freely available for download.

## Results

### Performances on Binding Affinity Benchmarks

In all tables for this section, we **bold** the best-performing model and *italicize* the second-best-performing model for each metric unless different CASTER-DTA formulations would be both the best and second-best, in which case we bold or italicize based on only the highest CASTER-DTA performance and instead underline other CASTER-DTA models that would have been the best or second-best had the other CASTER-DTA models not been present. For all datasets assessed, at least one of the four CASTER-DTA formulations performed best or second-best on every metric.

### Performance on Davis

On the Davis dataset (**Table 3**), the CASTER-DTA models outperform all of the comparison models on every metric with the exception of the MAE metric, where DGraphDTA performs best with an MAE of 0.229 while CASTER-DTA(1,2) has an MAE of 0.233.

**Table 3.**
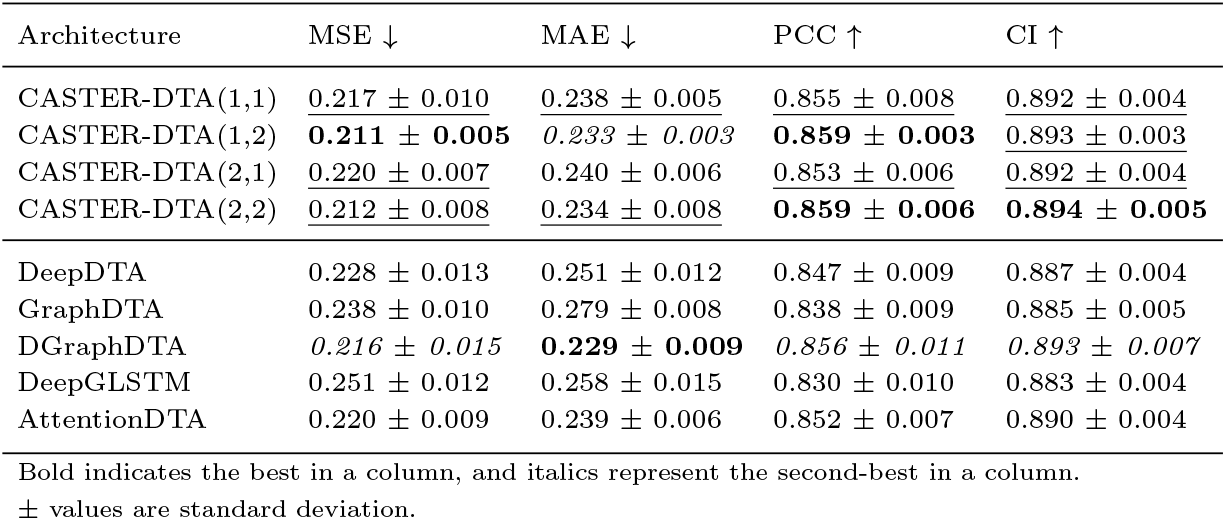
Performance on the Davis dataset

### Performance on KIBA

On the KIBA dataset (**Table 4**), the CASTER-DTA models outperform every comparison model except for DGraphDTA (MSE: 0.141; MAE 0.199; PCC 0.895; CI 0.897), where it nominally outperforms the CASTER-DTA models with the best one being CASTER-DTA(2,1) (MSE: 0.141; MAE 0.199; PCC 0.893; CI 0.896).

**Table 4.**
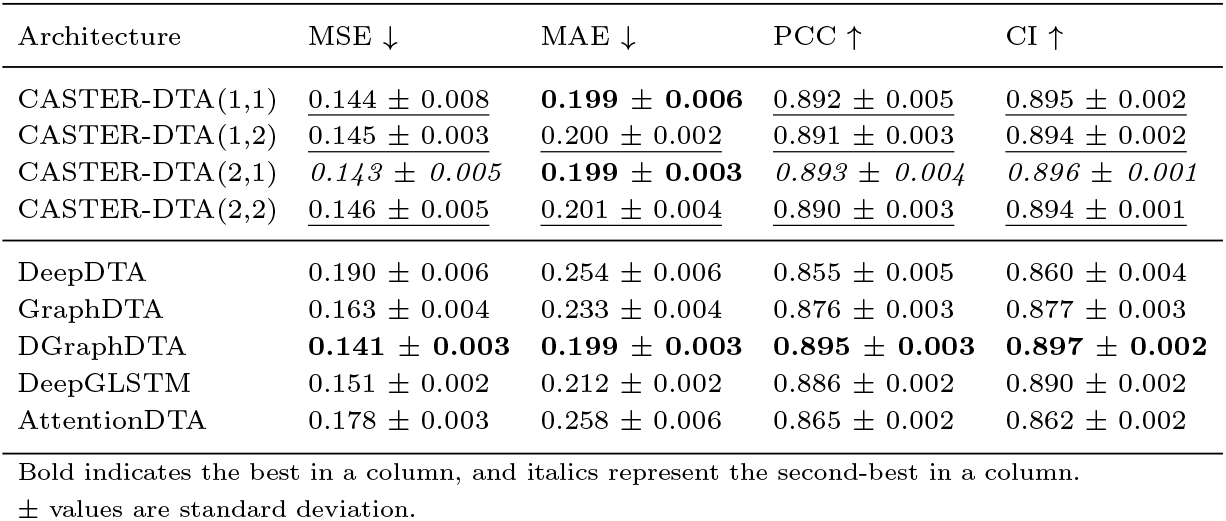
Performance on the KIBA dataset

| Architecture | MSE ↓ | MAE ↓ | PCC ↑ | CI ↑ |
| --- | --- | --- | --- | --- |
| CASTER-DTA(1,1) | <u>0.144 ± 0.008</u> | **0.199 ± 0.006** | <u>0.892 ± 0.005</u> | <u>0.895 ± 0.002</u> |
| CASTER-DTA(1,2) | <u>0.145 ± 0.003</u> | 0.200 ± 0.002 | 0.891 ± 0.003 | <u>0.894 ± 0.002</u> |
| CASTER-DTA(2,1) | *0.143 ± 0.005* | **0.199 ± 0.003** | *0.893 ± 0.004* | *0.896 ± 0.001* |
| CASTER-DTA(2,2) | <u>0.146 ± 0.005</u> | <u>0.201 ± 0.004</u> | <u>0.890 ± 0.003</u> | <u>0.894 ± 0.001</u> |
| DeepDTA | 0.190 ± 0.006 | 0.254 ± 0.006 | 0.855 ± 0.005 | 0.860 ± 0.004 |
| GraphDTA | 0.163 ± 0.004 | 0.233 ± 0.004 | 0.876 ± 0.003 | 0.877 ± 0.003 |
| DGraphDTA | **0.141 ± 0.003** | **0.199 ± 0.003** | **0.895 ± 0.003** | **0.897 ± 0.002** |
| DeepGLSTM | 0.151 ± 0.002 | 0.212 ± 0.002 | 0.886 ± 0.002 | 0.890 ± 0.002 |
| AttentionDTA | 0.178 ± 0.003 | 0.258 ± 0.006 | 0.865 ± 0.002 | 0.862 ± 0.002 |

Bold indicates the best in a column, and italics represent the second-best in a column. ± values are standard deviation.

### Performance on Metz and BindingDB

On both the Metz (**Table 5**) and BindingDB (**Table6**) datasets, CASTER-DTA exhibits consistently superior performance, with one or more of the CASTER-DTA models outperforming the available comparison models on every metric.

**Table 5.** Performance on the Metz dataset

| Architecture | MSE ↓ | MAE ↓ | PCC ↑ | CI ↑ |
| --- | --- | --- | --- | --- |
| CASTER-DTA(1,1) | <u>0.283 ± 0.005</u> | **0.392 ± 0.003** | **0.832 ± 0.004** | **0.815 ± 0.002** |
| CASTER-DTA(1,2) | <u>0.283 ± 0.007</u> | <u>0.393 ± 0.005</u> | **0.832 ± 0.006** | <u>0.814 ± 0.003</u> |
| CASTER-DTA(2,1) | <u>0.285 ± 0.011</u> | <u>0.393 ± 0.007</u> | <u>0.831 ± 0.008</u> | <u>0.814 ± 0.004</u> |
| CASTER-DTA(2,2) | **0.282 ± 0.006** | <u>0.394 ± 0.003</u> | **0.832 ± 0.005** | <u>0.814 ± 0.003</u> |
| DeepDTA | 0.317 ± 0.010 | 0.420 ± 0.006 | 0.810 ± 0.009 | 0.800 ± 0.004 |
| GraphDTA | 0.306 ± 0.007 | 0.414 ± 0.008 | 0.817 ± 0.006 | 0.806 ± 0.003 |
| DeepGLSTM | *0.292 ± 0.008* | *0.397 ± 0.005* | *0.826 ± 0.006* | *0.812 ± 0.003* |
| AttentionDTA | 0.301 ± 0.006 | 0.409 ± 0.004 | 0.821 ± 0.005 | 0.808 ± 0.005 |

Bold indicates the best in a column, and italics represent the second-best in a column. ± values are standard deviation.

**Table 6.** Performance on the BindingDB dataset

| Architecture | MSE ↓ | MAE ↓ | PCC ↑ | CI ↑ |
| --- | --- | --- | --- | --- |
| CASTER-DTA(1,1) | <u>0.729 ± 0.034</u> | **0.578 ± 0.011** | <u>0.853 ± 0.007</u> | <u>0.839 ± 0.004</u> |
| CASTER-DTA(1,2) | <u>0.732 ± 0.032</u> | <u>0.589 ± 0.010</u> | <u>0.853 ± 0.007</u> | <u>0.837 ± 0.003</u> |
| CASTER-DTA(2,1) | <u>0.741 ± 0.042</u> | <u>0.586 ± 0.013</u> | <u>0.851 ± 0.008</u> | <u>0.837 ± 0.004</u> |
| CASTER-DTA(2,2) | **0.717 ± 0.052** | **0.578 ± 0.022** | **0.856 ± 0.010** | **0.840 ± 0.006** |
| DeepDTA | 0.796 ± 0.029 | 0.627 ± 0.007 | 0.837 ± 0.006 | 0.828 ± 0.002 |
| GraphDTA | 0.740 ± 0.031 | *0.596 ± 0.006* | 0.850 ± 0.007 | 0.835 ± 0.003 |
| DeepGLSTM | *0.726 ± 0.031* | **0.578 ± 0.010** | *0.852 ± 0.006* | *0.838 ± 0.003* |
| AttentionDTA | 0.777 ± 0.034 | 0.620 ± 0.011 | 0.841 ± 0.007 | 0.829 ± 0.003 |

Bold indicates the best in a column, and italics represent the second-best in a column. ± values are standard deviation.

**Table 7.**
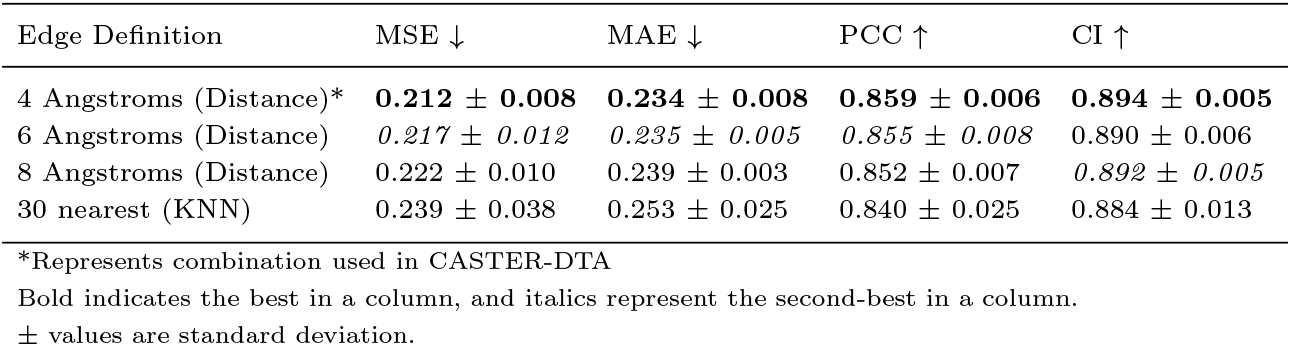
Performance of CASTER-DTA with Different Edge Definitions (on Davis)

| Edge Definition | MSE ↓ | MAE ↓ | PCC ↑ | CI ↑ |
| --- | --- | --- | --- | --- |
| 4 Angstroms (Distance)* | **0.212 ± 0.008** | **0.234 ± 0.008** | **0.859 ± 0.006** | **0.894 ± 0.005** |
| 6 Angstroms (Distance) | *0.217 ± 0.012* | *0.235 ± 0.005* | *0.855 ± 0.008* | 0.890 ± 0.006 |
| 8 Angstroms (Distance) | 0.222 ± 0.010 | 0.239 ± 0.003 | 0.852 ± 0.007 | *0.892 ± 0.005* |
| 30 nearest (KNN) | 0.239 ± 0.038 | 0.253 ± 0.025 | 0.840 ± 0.025 | 0.884 ± 0.013 |

\*Represents combination used in CASTER-DTA
Bold indicates the best in a column, and italics represent the second-best in a column.
± values are standard deviation.

### Ablation Studies

#### Edge Definition Ablation Performance

As seen in **Table 7**, the edge definition of 4 Angstroms performed the best of all of the edge definitions on all metrics across MSE, MAE, PCC, and CI. Notably, all definitions including the 30 nearest neighbors definition performed well, with many of these definitions still outperforming most of the current state-of-the-art methods on the Davis dataset (described in **Table 3**).

As the 4 Angstrom edge definition offered significant speedup in training and less memory usage due to needing creating much sparser protein graphs (generally requiring approximately 1/5 of the training time compared to the 30 nearest neighbors definition), we used it as the basis for the protein graphs in CASTER-DTA.

#### GNN Architecture Ablation Performance

As seen in **Table 8**, the architecture using the GVP-GNN for proteins and GINE for molecules achieved the best performance of all of the architectures evaluated, with other architectures using the GVP-GNN for the protein showing MSEs in the range of 0.215-0.219 and the architecture replacing the GVP-GNN with GAT showing an MSE of 0.231. PCC and CI show similar trends, with the GVP-GNN + GINE combination as used in CASTER-DTA outperforming all other combinations, and the GVP-GNN + GINE combination was best in the MAE metric with an MAE of 0.234 compared to the second-best with an MAE of 0.236 in the GVP-GNN + AttentiveFP combination. We ultimately used the GVP-GNN + GINE combination in CASTER-DTA as it exhibited superior performance and considerable stability in training across hyperparameters, requiring less tuning and training time.

**Table 8.** Ablation Study of Various GNN Architectures (on Davis)

| Protein GNN | Molecule GNN | MSE ↓ | MAE ↓ | PCC ↑ | CI ↑ |
| --- | --- | --- | --- | --- | --- |
| GVP-GNN* | GINE* | **0.212 ± 0.008** | **0.234 ± 0.008** | **0.859 ± 0.006** | **0.894 ± 0.005** |
| GATv2 | GINE | 0.231 ± 0.011 | 0.244 ± 0.005 | 0.845 ± 0.008 | 0.886 ± 0.005 |
| GVP-GNN | GIN | 0.219 ± 0.008 | 0.237 ± 0.005 | 0.854 ± 0.007 | 0.891 ± 0.003 |
| GVP-GNN | GATv2 | *0.215 ± 0.011* | 0.238 ± 0.008 | *0.857 ± 0.008* | **0.894 ± 0.006** |
| GVP-GNN | AttentiveFP | 0.216 ± 0.010 | *0.236 ± 0.005* | 0.856 ± 0.007 | *0.893 ± 0.004* |

\*Represents combination used in CASTER-DTA
Bold indicates the best in a column, and italics represent the second-best in a column.
± values are standard deviation.

### Interpretability Analyses

#### BioLIP Binding Residues

The distribution of the differences in attention between binding and non-binding residues can be seen in **Figure 2**. Based on the paired t-test on the binding versus non-binding residue attention scores, we rejected the null hypothesis that the difference in attention scores was zero (*t* = 6.5502, *p* =9.472 *×* 10^*−*11^, *df* = 935, *µ*_diff_ = 0.0011), indicating that CASTER-DTA preferentially places attention on putative binding residues as defined by the BioLIP dataset. For visual validation, we also plot the attention scores over all residues in scatter plots for select proteins where the difference in attention scores was the largest, as well as the average attention score over all residues and the average attention score over binding residues, in **Figure3**.

**Fig. 2.**
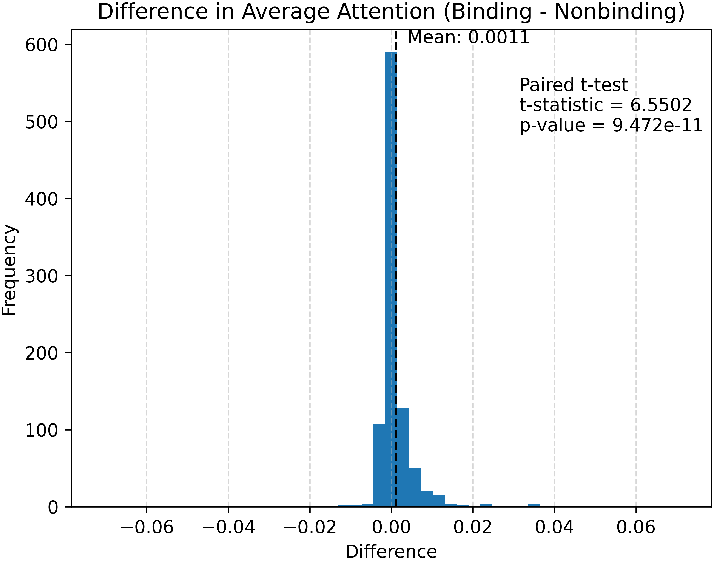
Distribution of differences in attention scores over binding versus nonbinding residues in each protein, with paired t-test for testing the hypothesis that there was no difference listed. The null hypothesis was rejected with *t* = 6.5502, *p* = 9.472 *×* 10^*−*11^, *df* = 935, *µ*_diff_ = 0.0011.

**Fig. 3.**
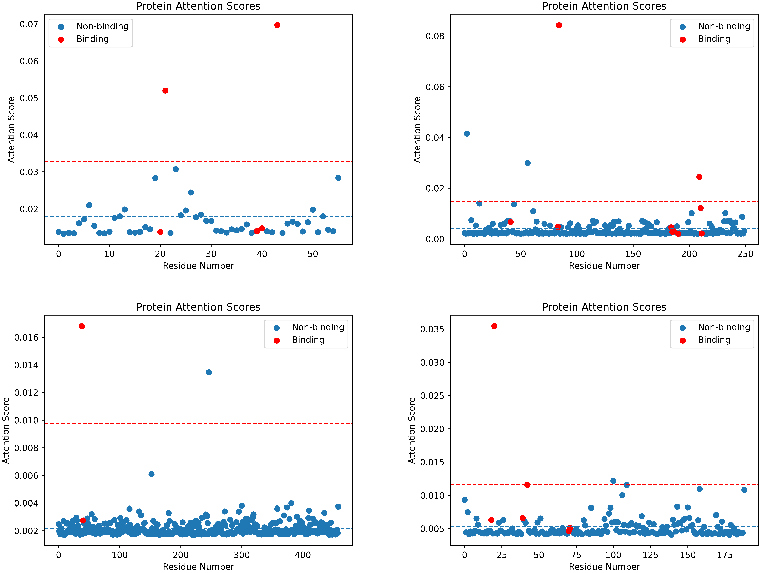
Scatter plot of residue number (0-indexed) against attention scores for each residue; BioLIP binding residues are colored in red while nonbinding residues are colored in blue. The average attention score is represented by the blue dashed line, while the average binding residue attention score is represented by the red dashed line. Top left: protein Q07654 bound to BioLIP ligand GAL; top right: protein P04070 bound to BioLIP ligand 0G6; bottom left: protein O95470 bound to BioLIP ligand P04; bottom right: protein P19440 bound to BioLIP ligand GLU.

Some of the residues with higher attention scores are not some of the putative residues that we would expect or that some of the binding residues are given attention scores more in line with the nonbinding residues, possibly because BioLIP defines putative binding residues based on proximity, and residues farther away from a ligand can still be important in determining binding affinity in a variety of ways (for example, by limiting how much regions of the protein can move to conform to a ligand).

### PharmGKB Mutation Impact

A summary of the results produced by CASTER-DTA can be seen in **Table 9**, with the four tested mutations all showing a decrease in binding affinity to the respective drugs (requiring more drug for the alternate protein than the original, reference protein). A visualization of the four mutations and their differences can also be found in **AppendixD**.

**Table 9.** PharmGKB Mutation Assessment Results

| Variant ID | Gene | AA Change | Attention Near? | Ratio (Alt./Ref.) | Matches Lit.? |
| --- | --- | --- | --- | --- | --- |
| rs61742245 | VKORC1 | Asp36Tyr | Yes (35) | 1.65 | Yes [26] |
| rs1800462 | TPMT | Ala80Pro | Yes (80) | 1.47 | Yes [27] |
| rs4149056 | SLCO1B1 | Val174Ala | Yes (166) | 2.04 | Yes [28] |
| rs2306283 | SLCO1B1 | Asn130Asp | Yes (134, 138) | 2.05 | Unclear [29] |

A ratio $>1$ means more drug is needed to bind to the alternate protein (decreased affinity).

A similar pattern to the BioLIP analysis can be seen in this analysis with the residues highlighted by CASTER-DTA and how their attention scores change from normal protein to mutant protein: some of the highlighted residues are those physically *near* or at the mutated residue, even when that residue itself may not be the location where the drug binds to the protein.

This can be seen by comparing the two latter mutations (rs4149056 and rs2306283) which are both on the same protein (NP 006437.3, SLCO1B1) and affect the same ligand (atorvastatin) but where the mutations themselves in different locations. Here, the expected binding pocket of the ligand is expected to be the same; however, despite the mutations being in different locations, CASTER-DTA shows that some residues near these mutations as well as some farther away exhibit a change in attention, which may indicate a dual focus on residues directly bind to the residue as well as residues that mediate the binding process. The fact that some of the top residues with changed attention are consistent between these two variants while others (particularly those around the mutated residues) are different lends credence to this interpretation and to the utility of CASTER-DTA in assessing mutations.

Additionally, we see that the change in binding affinity as predicted by CASTER-DTA matches experimental observations, with all mutations in question generally reducing predicted affinity. This provides further evidence that CASTER-DTA can effectively distinguish between proteins that are otherwise very similar structurally and characterize protein-coding mutations in the context of binding.

### Drug Repurposing

#### Validation through Protein Case Studies

The top three drug candidates for repurposing (ranked by binding affinity) for each protein can be found in **Table 10**. We report for each protein-drug pair whether there is evidence that the drug can be used to target the protein in some capacity. For some of the drugs found in this manner, there exists evidence that they may be useful in targeting their respective protein, while many others are novel.

**Table 10.** Top Binding-Affinity Drugs for each Protein Assessed

| Protein | Drug | Prior Evidence? |
| --- | --- | --- |
| Amyloid-beta peptide | Adapalene | Yes [30] |
|  | Brinzolamide | No |
|  | Lopinavir | Some [31] |
| Amyloid precursor protein | Indigotindisulfonic acid | No |
|  | Tenoxicam | No |
|  | Fluorescein | No |
| Acetylcholinesterase | Repotrectinib | No |
|  | Sirolimus | Some [32] |
|  | Pimecrolimus | No |
| Tau | Temsirolimus | No |
|  | Sirolimus | No |
|  | Afamelanotide | No |
| H5 Hemagglutinin | Dorzolamide | Some [33] |
|  | Lopinavir | No |
|  | Brinzolamide | Some [34] |
| H5N1 Nucleoprotein | Methotrexate | Some [35] |
|  | Brinzolamide | Some [34] |
|  | Pralatrexate | Some [35] |
| N1 Neuraminidase | Lopinavir | No |
|  | Quinestrol | No |
|  | Chloramphenicol palmitate | No |

Some of the highest-binding-affinity drugs to Amyloid-beta peptide include adapalene and lopinavir, which have independently been put forth as possible drug candidates. Adapalene has been found to inhibit amyloid-beta aggregation in vitro[30]. Lopinavir was shown to reduce amyloid-beta levels in vitro through an undetermined mechanism but failed to show effects in vivo due to likely-poor penetration into the nervous system[31].

Additionally, the affinity of various known drugs for Acetylcholinesterase and N1 Neuraminidase are found in **Table 11**. All of the drugs have a higher binding affinity than the average of all drugs against that protein (z-score *>* 0), with some being in the top percentile of all possible binding drugs such as Donepezil (with a z-score of 1.60 or close to 10th percentile).

**Table 11.**
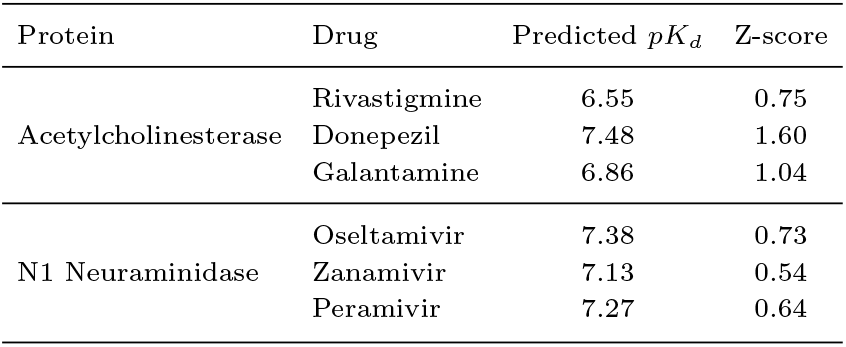
Affinity of Known Drugs for Acetylcholinesterase and N1 Neuraminidase

| Protein | Drug | Predicted $pK_d$ | Z-score |
| --- | --- | --- | --- |
| Acetylcholinesterase | Rivastigmine | 6.55 | 0.75 |
|  | Donepezil | 7.48 | 1.60 |
|  | Galantamine | 6.86 | 1.04 |
| N1 Neuraminidase | Oseltamivir | 7.38 | 0.73 |
|  | Zanamivir | 7.13 | 0.54 |
|  | Peramivir | 7.27 | 0.64 |

### Resource of Millions of Binding Affinities across the Entire Human Proteome

We make freely available for download a prediction of binding affinities for the entire human proteome against every FDA-approved drug identified as of March 2025. This resource comprises 37,776,465 binding affinities for 1615 FDA-approved drugs against 23,391 predicted structures (20,504 unique proteins, with some proteins having multiple fragments). Proteins are identified by their UniProt ID and sequence (as well as the fragment ID for larger proteins) and molecules are identified by their ZINC ID and SMILES. For ease of access and compression, the data is made available in the Parquet file format (which allows for high compression of the data). One can load the entire dataframe using the Python Pandas pandas.read_parquet function on this file.

### Web Server

We make it possible for researchers to run CASTER-DTA on a limited set of proteins and drugs of their own choosing by creating a web server where users can upload their own protein structure in the PDB format and enter a drug SMILES to be analyzed by CASTER-DTA. The web server then runs the same pretrained CASTER(2,2) DTA model used in all of the downstream tasks above and provides to the user several results for their own utility: the binding affinity of the protein-drug pair, the attention scores of the protein residues, and the attention scores of the drug atoms. It also visualizes the protein attention overlaid on the 3D structure for the user, allowing them to identify residues important for prediction. The web server can be found at https://caster.ritchielab.org.

## Discussion

CASTER-DTA in all of its iterations [(1,1), (1,2), (2,1), and (2,2)] exhibits remarkably consistent performance that exceeds or matches those of other models across multiple datasets; on every dataset tested, all four CASTER-DTA models tested were at least second-best on every metric. On the Davis, Metz, and BindingDB datasets, one or more CASTER-DTA models outperformed every comparison architecture, and on the KIBA dataset, all of the CASTER-DTA models were second only to DGraphDTA but otherwise achieved performance similar to it on every metric.

Ultimately, CASTER-DTA is able to perform well in predicting binding affinity with no external information beyond the structural information already contained within a protein other than fundamental amino acid properties (such as molecular weight, hydrophobicity, and others), which are fixed values for each residue type across proteins. This offers it significant flexibility, and it is able to perform well on all four tested datasets, with the CASTER-DTA models generally performing best or second-best on every dataset regardless of convolution counts, indicating CASTER-DTA’s robustness to architectural settings and to different dataset domains in protein and molecule distributions.

Additionally, we find through our ablation studies evidence that the SE(3)-equivariant GVP-GNN is able to effectively make use of protein 3D structural information to generate equivariant representations and that these representations improve the performance of CASTER-DTA compared to other baselines. Specifically, when we replace the protein GVP-GNN in CASTER-DTA with a GATv2 architecture (which does not consider the 3D protein information in an equivariant manner), we find that performance is noticeably reduced, with the MSE increasing from 0.212 to 0.231 (**Table 8**). Furthermore, when using other molecular GNN architectures such as GIN, GATv2, and AttentiveFP, we find that they maintain performance when run alongside the GVP-GNN, even continuing to outperform some of the existing state-of-the-art methods. This indicates that the equivariant protein representations produced by the GVP-GNN are highly adaptable and can work alongside a variety of molecular representations.

From the BioLIP and PharmGKB analyses, CASTER-DTA is able to highlight residues that we would expect it to, indicating its value for downstream analyses that need interpretability. For BioLIP specifically, we see that the average attention given to binding residues is higher than the average attention score given to non-binding residues for a variety of protein-ligand complexes, indicating that the attention mechanism does indeed generally focus on residues that we would expect it to when determining binding affinity (**Figure 2**). For PharmGKB, we see that CASTER-DTA generally identifies residues near to or at the location of mutations known to affect protein pharmacokinetics (**Table 9**). Given CASTER-DTA’s ability to highlight relevant residues for binding, it may have the potential to be used to design drugs that target specific residues of a protein.

When used to rank drugs as possible candidates for binding to specific proteins, CASTER-DTA is able to find several drugs with prior evidence supporting their potential use, including lopinavir and adapalene for Amyloid-beta peptide and sirolimus for Acetylcholinesterase (**Table 10**). CASTER-DTA also assigns higher-than-average binding affinities to drugs known to target certain proteins such as Acetylcholinesterase and N1 Neuraminidase (**Table 11**) compared to the landscape of all FDA-approved drugs. Taken together, this shows that CASTER-DTA can either rank an existing list of drug candidates or to generate a starting list of promising candidates *de novo* based on binding affinity alone. However, CASTER-DTA only predicts binding affinity and so cannot account for other considerations such as blood-brain barrier penetration or tissue distribution.

## Conclusion

In summary, we present a model known as CASTER-DTA (Cross-Attention with Structural Target Equivariant Representations for Drug-Target Affinity) that makes use of graph neural networks to process 3D protein structural information in an equivariant manner to predict drug-target affinity. We show that CASTER-DTA is able to perform consistently well across datasets, outperforming many of the existing state-of-the-art methods on several datasets. We additionally show that CASTER-DTA’s cross-attention module allows one to introspect what components of a protein are contributing to the prediction and that CASTER-DTA consistently focuses on residues that are putatively involved in binding as well as adjusts its attention between a normal and mutant protein appropriately, with it being sensitive to changes in protein sequence in pharmacogenetic applications. Overall, CASTER-DTA is a useful tool for predicting binding affinity with great potential in drug development and discovery applications.

## AlphaFold2/ColabFold Parameters

Parameters that were used to fold any proteins not already present in the Protein DataBank or the AlphaFold2 database can be found in **Table A1**.

**Table A1.**
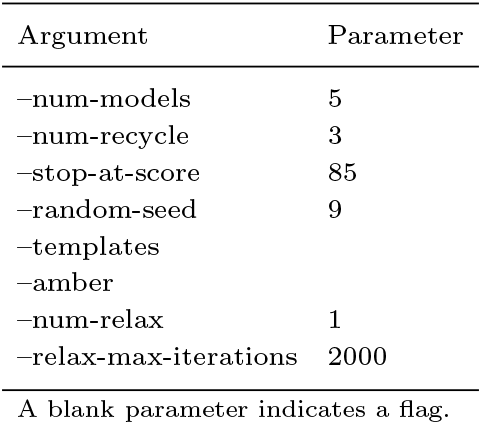
ColabFold parameters

## Graph Preparation

A figure that provides some details on the process of constructing protein and molecular graphs can be found in **Figure B1**, with additional details on the features in each graph type in the subsections below.

**Fig. B1.**
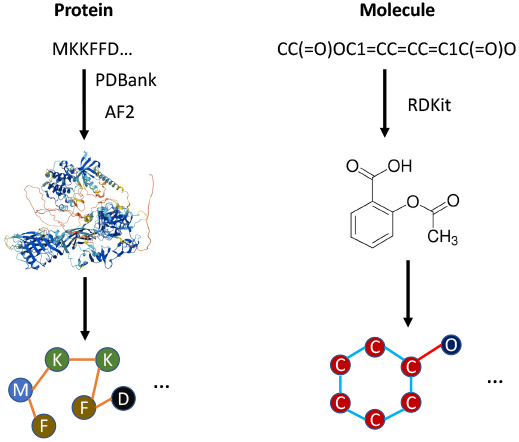
Avisualization of the process of generating protein and molecular graphs, starting from protein sequence and molecular SMILES strings. Protein structures are acquired from PDBank or folded with AlphaFold2 and molecule SMILES are converted to graphs using RDKit.

### Protein Graph Features

#### Node Features

Each node (residue) has 17 scalar features and 3 vector features:

- Sin and cos of the three dihedral angles computed from the residue’s central carbon to its bonded atoms. (6 scalars)
- Various amino acid properties, including weights, pK and pI, hydrophobicity, and binary features of aliphaticity, aromaticity, whether the residue is acidic or basic, and whether the residue is polar neutral. (11 scalars)
- A unit vector pointing from the residue’s central carbon towards the sidechain, computed by normalizing from tetrahedral geometry. (1 vector)
- One unit vector each pointing from the residue’s central carbon to the central carbons of the previous and next residues in sequence. The first and last residues have zero-vectors for their backward and forward vectors, respectively. (2 vectors)

### Edge Features

Each edge (based on distance) has 32 scalar features and 1 vector feature:

- Gaussian RBF encodings of the distance between the residues. (16 scalars)
- Sinusoidal positional encodings of the difference in residue sequence indices. (16 scalars)
- Unit direction vector pointing from the source residue’s central carbon to the destination residue’s central carbon. (1 vector)

### Molecule Graph Features

#### Node Features

Each node (atom) has 41 total features:

- One-hot encodings of various properties, including: chirality, hybridization, number of bound hydrogens, degree, valence, formal charge, and number of electron radicals. (38 features)
- Binary features for whether the atom is in a ring and is aromatic. (2 features)
- Computed Gasteiger partial charge, if available (if not, set to 0.0). (1 feature)

### Edge Features

Each edge (bond) has 9 features (self-loops have all features set to 0):

- Bond stereoconfiguration. (7 features)
- Binary features for whether the bond is conjugated and in a ring. (2 features)

## CASTER-DTA Architecture and Comparison Method Details

The CASTER-DTA architecture is described in detail in **Figure C1**. Information about the implementations of other benchmark drug-target affinity methods can be found in **Table C1**. All methods were implemented in PyTorch (for DeepDTA, this required reimplementation).

**Table C1.** Parameters and Settings for Benchmark Methods

| Method | Code source | LR | Epochs | Batch |
| --- | --- | --- | --- | --- |
| DeepDTA | Reimplemented | 0.001 | 100 | 256 |
| GraphDTA | Original code | 0.0005 | 1000 | 128 |
| DGraphDTA | Original code | 0.001 | 2000 | 512 |
| DeepGLSTM | Original code | 0.0005 | 1000 | 128 |
| AttentionDTA | Original code | 0.0001 | 100 | 512 |

## PharmGKB Mutations

A visualization of the four mutations assessed can be found in **Figures D1, D2, D3, and D4**. The top left of each figure represents the reference protein and its corresponding attention log-ratios, the top right represents the mutant protein and its attention log-ratios, the bottom left overlays a ratio of the attention log-ratios at each residue on top of the reference protein, and the bottom right shows numerically the top 10 residues with changes in attention from reference to mutant overlaid on where the protein would be in 3D space. Yellow on the protein indicates that the attention is close to the average (or that the ratio between reference and mutant is close to 1), red indicates a decrease in attention (compared to the average

**Fig. C1.**
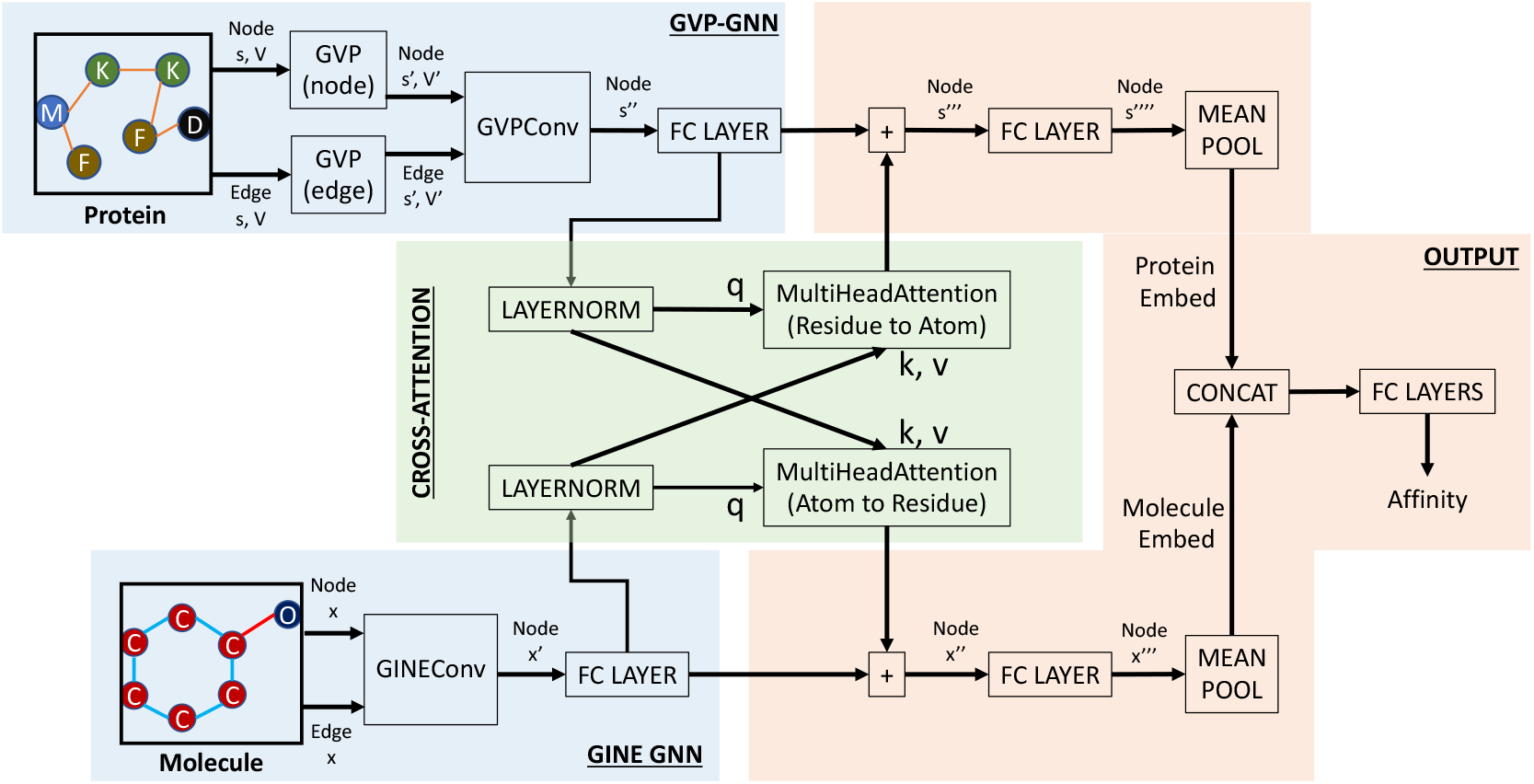
Avisualization of the architecture used to predict drug-target affinity from protein and molecule graphs. GVP = Geometric Vector Perceptron; GINE = Graph Isomorphism Network with Edge Enhancement.

**Fig. D1.**
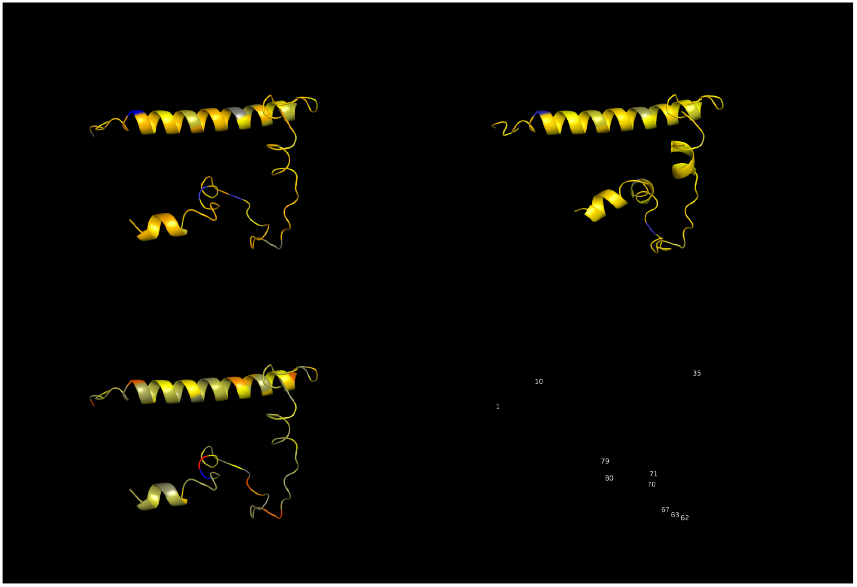
For rs61742245 (NP 996560.1:p.Asp36Tyr). A visualization of the attention or differences thereof on the protein.or from reference to mutant), and blue indicates an increase in attention.

**Fig. D2.**
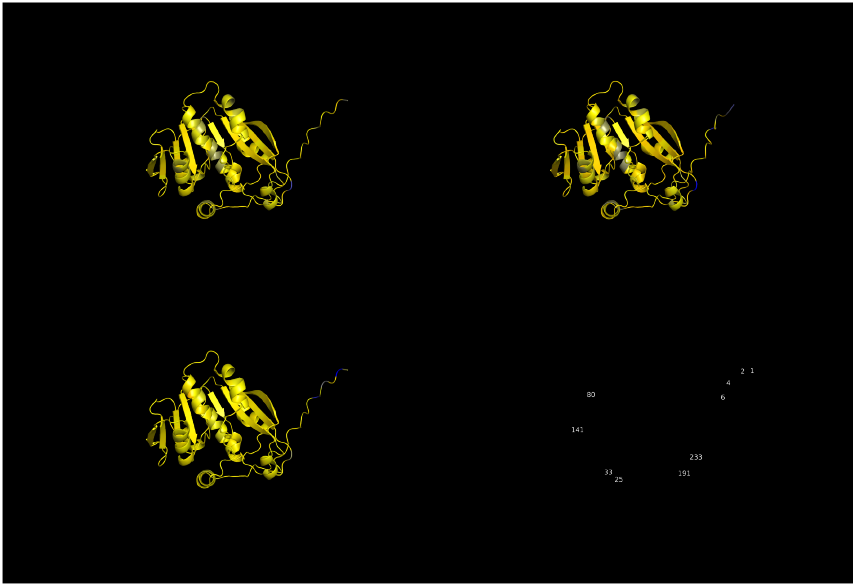
For rs1800462 (NP 000358.1:p.Ala80Pro). A visualization of the attention or differences thereof on the protein.

## Assessed Proteins for Drug Repurposing

A table with the details of each protein that was assessed for drug repurposing as well as the PDBank structure used for each protein can be found in **Table E1**.

## Data and Code Availability

The underlying code and the datasets used for this study can be accessed via this link: https://github.com/rachitk/caster-dta. The datasets are either freely available and can be redistributed (Davis, KIBA, Metz) or are freely available for download from the original source (BindingDB).

The proteome-wide binding affinity data will be made available through a data repository upon publication. A web server for CASTER-DTA, as described in the paper, is available at: https://caster.ritchielab.org.

**Fig. D3.**
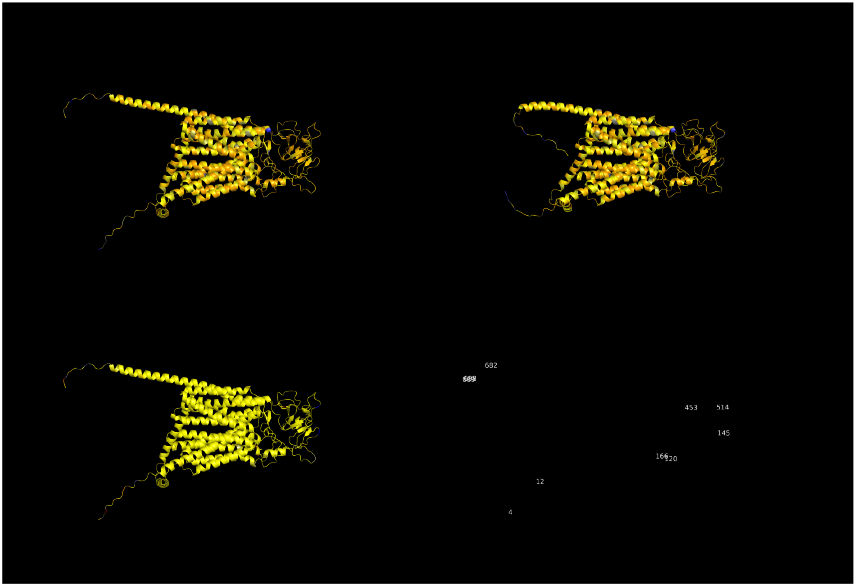
For rs4149056 (NP 006437.3:p.Val174Ala). A visualization of the attention or differences thereof on the protein.

**Fig. D4.**
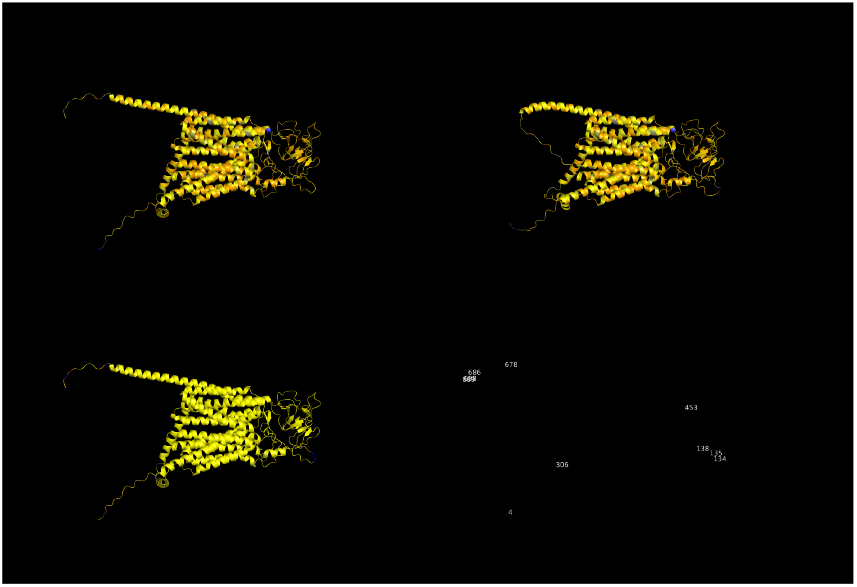
For rs2306283 (NP 006437.3:p.Asn130Asp). A visualization of the attention or differences thereof on the protein.

**Table E1.** Proteins Assessed for Drug Repurposing

| Protein | Disease | PDB ID | Description |
| --- | --- | --- | --- |
| Amyloid-beta peptide | Alzheimer's disease | 6SZF | Tends to aggregate into plaques thought to contribute to AD |
| Amyloid precursor protein | Alzheimer's disease | AF_AFP05067F1 | Protein that is cleaved to generate A-beta. |
| Acetylcholinesterase | Alzheimer's disease | AF_AFP04058F1 | Target to try to alleviate symptoms of AD; may be involved in pathogenesis |
| Tau | Alzheimer's disease | AF_AFP27348F1 | Aggregates into insoluble filaments thought to cause Alzheimer's disease |
| H5 Hemagglutinin | H5N1 influenza | 4MHI | Protein that binds to receptors on target cells to enable infection |
| H5N1 Nucleoprotein | H5N1 influenza | 2Q06 | Performs many roles in binding of virus to host and allowing for viral replication |
| N1 Neuraminidase | H5N1 influenza | 2HTY | Helps virus escape host cells by cleaving sialic acid on the surface of the cell |

## Author Contributions

RK conceived the idea for this study, wrote the code for the analyses performed in this paper, wrote the initial draft of the paper, and edited the final draft of the paper. JDR provided supervision, code support regarding the graph neural network models, and edited the final draft of the paper. MDR provided supervision, funding support, advice on analyses to perform, and edited the final draft of the paper.

## Acknowledgments

RK was supported by the National Human Genome Research Institute of the National Institutes of Health (T32HG000046). JDR was supported by the National Library of Medicine of the National Institutes of Health (R00LM013646). MDR was supported by the National Institute on Aging of the National Institutes of Health (U01AG066833). The funders played no role in study design, data collection, analysis and interpretation of data, or the writing of this manuscript.

**Rachit Kumar**. Rachit is a current MD-PhD student at the Perelman School of Medicine at the University of Pennsylvania, and he recently completed his PhD in Genomics and Computational Biology. His dissertation research was on the use of graph neural networks for interpretable biomedical data analysis.

